# Dynamic Tuning of Genetic Circuit Gain via Controllable Plasmid Copy Number

**DOI:** 10.1101/2025.08.13.670071

**Authors:** Haruno Namura, Yutaka Hori

**Author notes:** Email: {, }.

## Abstract

Plasmid copy number directly influences gene expression strength in synthetic genetic circuits and serves as a tunable parameter for adjusting circuit gain and output levels. Building upon recent progress in copy number control, this report presents a genetic circuit that enables dynamic regulation of circuit gain through the regulation of plasmid copy numbers. To illustrate the underlying concept, we first developed a mathematical model describing the relationship between plasmid copy number and circuit gain and simulated the circuit dynamics to examine how gain responds to copy number changes. We then constructed the circuit and experimentally confirmed that plasmid copy number, *i*.*e*., circuit gain, can be externally controlled in *E. coli*. In particular, we demonstrate communication-based gain control using AHL-mediated quorum sensing from co-cultured cells. These results support the feasibility of plasmid copy number control as a practical and predictable strategy for tuning synthetic gene circuit behavior.

## 1 Introduction

Optimizing the volume of protein synthesis, the output of genetic circuits, is crucial for achieving desired cellular responses. To this end, it is often necessary to adjust how sensitively a genetic circuit responds to input signals such as inducers, environmental cues, or transcription factors generated by upstream circuits. This sensitivity can be quantitatively characterized by the *gain* of the circuit, which is a concept borrowed from control engineering, defined as the ratio between changes in output and the corresponding changes in input.^1^ Traditionally, the gain adjustment in synthetic biocircuit has relied on recombining genetic parts such as promoters and ribosome binding sites, thereby modulating the transcriptional and translational rates.^3–6^ However, the nonlinear nature of these biochemical processes makes it difficult to fine-tune the resulting input-output relationship, and achieving desired output levels often requires laborious iterative exploration.

As an alternative tuning knob, recent studies have explored the use of dynamically controllable plasmid copy number to modulate gene expression levels.^7–10^ These studies demonstrated that altering the number of plasmids within a cell can significantly affect protein synthesis levels, as a higher plasmid copy number typically leads to increased transcriptional activity due to a greater number of gene templates. In particular, this strategy offers orthogonality to traditional transcriptional or translational modifications, allowing circuit designers to independently adjust gain without re-engineering core regulatory elements. The control of plasmid copy number was previously leveraged to build a genetic circuit for population-level synchronization through regulated plasmid cycling^11^ and pathway amplification using inducible control of plasmid replication.^12^

Building upon these works, this report presents a genetic circuit that enables external control of plasmid copy number. While our approach shares the same molecular mechanism as in the previous study,^9^ it introduces a distinct circuit topology designed for gain adjustment in synthetic genetic circuits using widely used regulatory proteins and inducers. Our design of the circuit avoids introducing negative feedback in the expression of the replication initiator protein, making the system’s response to external input more predictable and easier to characterize. To illustrate the concept, we first formulated the proposed circuit as an input-output system with variable gain and simulated its dynamics to confirm that the gain increases linearly with the plasmid copy number. Then, to validate the design, we constructed the circuit and experimentally confirmed that the plasmid copy number, or the circuit gain, can be dynamically regulated by external inputs. In particular, we showed that copy number can be modulated via AHL-based quorum sensing signals secreted from co-cultured *E. coli* cells, enabling communication-driven gain control by neighboring cells. This highlights the potential of copy number modulation as a versatile strategy for implementing tunable and cooperative behaviors in synthetic biology.

## 2 Results

### 2.1 Mathematical model of biocircuits with variable copy number

In this section, we introduce a conceptual input-output model of the biocircuit with variable plasmid copy number and demonstrate through computational simulations that the circuit gain can be modulated by the copy number of the circuit plasmid.

We consider a biocircuit consisting of a plasmid with variable copy number that encodes the gene for the output protein, Protein_x_, whose transcription is regulated by the upstream regulator, Protein_u_. The copy number of the plasmid, which we hereafter denote by Plasmid_k_, is regulated by a protein-mediated replication mechanism, in which an initiator protein Protein_d_ binds to the specific origin of replication (ori) on the plasmid to facilitate the initiation of plasmid replication. Thus, Protein_d_ is a regulator of the copy number of Plasmid_k_. An example of such protein-mediated replication control is the RepA protein-based regulation in the pSC101 plasmid, where the initiator protein RepA specifically binds to the origin of replication to control its replication frequency.^13^ In such systems, modulation of RepA concentration results in the difference in plasmid copy number, providing a mechanism to dynamically regulate gene dosage in synthetic circuits.^9, 14^

A conceptual block diagram showing an input-output relation of the biocircuit is illustrated in Fig. 1. In the figure, *u*(*t*) and *x*(*t*) represent the concentrations of the input protein, Protein_u_, and the output protein, Protein_x_, respectively, and *d*(*t*) represents the effective rate of change in plasmid concentration per unit time, accounting for both replication and dilution effects due to cell division. The plasmid concentration modulates the overall input-output responsiveness of the circuit, which we refer to conceptually as the gain of the circuit. This gain reflects how strongly changes in the input signal affect the output response and can be viewed as the ratio *x*(*t*)*/u*(*t*) at steady state.

**Figure 1:**
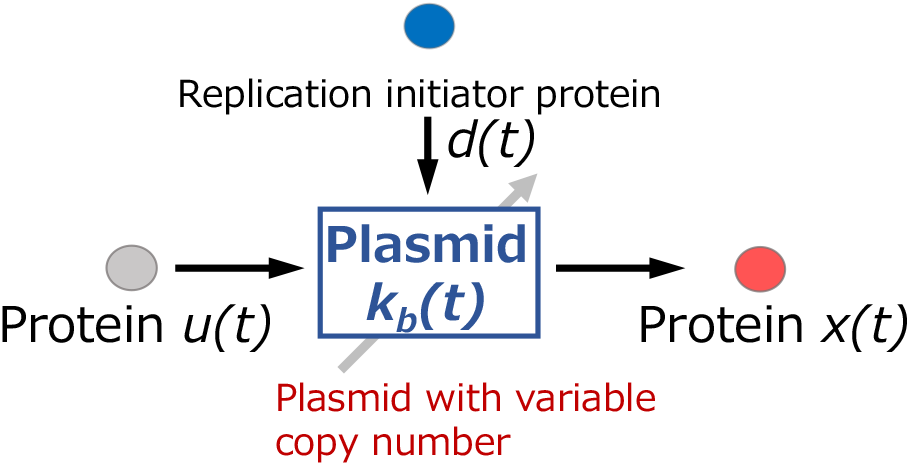
A block diagram of genetic circuit with variable plasmid copy number

More specifically, the binding-dissociation reaction between the input protein, Protein_u_, and Plasmid_k_, as well as the synthesis and degradation reactions of the input protein, Protein_u_ and output protein, Protein_x_, are modeled by the following formula.

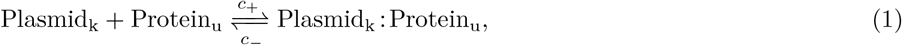

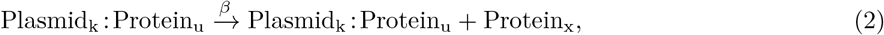

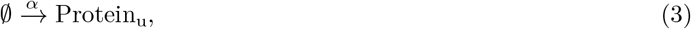

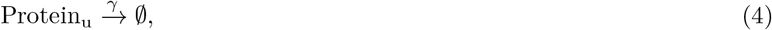

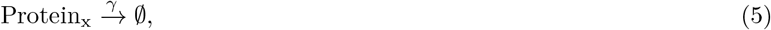

where Plasmid_k_ : Protein_u_ represents a complex of Protein_u_ and Plasmid_k_ (Protein_u_ bound to the promoter on the plasmid Plasmid_k_), and the input Protein_u_ is assumed to be expressed constitutively. The constants *c*_+_ and *c*_−_ are the rates of binding or dissociation, respectively, *α* and *β* represent the production rate of each protein, and *γ* is the degradation (dilution) rate of proteins.

Using the law of mass action, we approximate the dynamics of the reactions with continuous concentrations, including plasmid copy number and the effects of dilution due to cell growth. The resulting differential equations describe the temporal evolution of protein and plasmid concentrations.

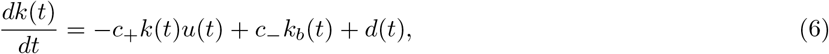

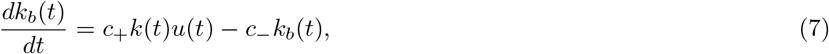

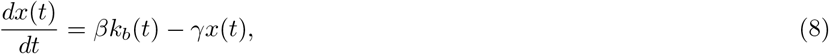

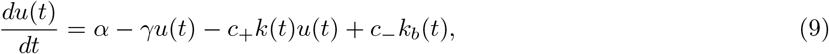

where *k*(*t*) and *k*_*b*_(*t*) are the concentrations of the free plasmid, Plasmid_k_, and the bound plasmid, Plasmid_k_ : Protein_u_, respectively. Thus, the total concentration of the plasmid is *k*(*t*) + *k*_*b*_(*t*), and the rate of its increase is *d*(*t*) as seen from (6) and (7).

The rate *d*(*t*) is dependent on the concentration of the replication initiator protein, Protein_d_, thereby the plasmid copy number can be modulated by tuning the expression level of Protein_d_. Eliminating *k*(*t*), eq. (7) can be expressed using the change rate of plasmid *d*(*t*) as

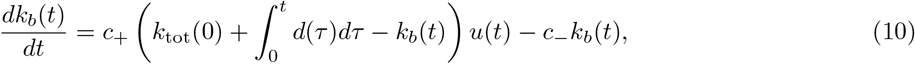

where *k*_tot_(0) is the initial copy of plasmid. This equation shows how the effective change rate of copy number *d*(*t*) acts as a parameter to tune the system’s sensitivity to the input *u*(*t*), thereby modulating the output *x*(*t*) through eq. (8). As intuitively expected, the output *x*(*t*) increases linearly with the total plasmid copy number at steady state.

### 2.2 Numerical demonstration of gain tuning via control of plasmid copy number

To numerically observe the conceptual framework, we simulated the dynamics of the system based on eqs. (6)–(9) while varying *d*(*t*). Specifically, the following two scenarios were considered: (i) the total plasmid concentration *k*_tot_(*t*) = *k*(*t*) + *k*_*b*_(*t*) was kept constant, *i*.*e*., *d*(*t*) = 0, at high (300 nM) and low (100 nM) level, and (ii) the total plasmid concentration was ramped at a constant rate *d*(*t*) = 1 for 30 ≤ *t* ≤ 230. The parameters used in this simulation are shown in Table 1. The binding rate between Plasmid_k_ and Protein_u_, *c*_+_, was set close to the diffusion limit, assuming a high-affinity interaction, and the dissociation rate *c* was determined from the assumed equilibrium dissociation constant *K*_*d*_ := *c*_−_ */c*_+_ 1.0.^15^ The rate of Protein_x_ production includes contributions from transcription, translation, and protein folding time, with values taken from the literature.^16^ The degradation rate of proteins was determined assuming a cellular half-life of 20 minutes.

**Table 1:** Parameters for the simulations.

| Description | Symbol | Value | Unit |
| --- | --- | --- | --- |
| Dissociation of Plasmid<sub>k</sub>:Protein<sub>u</sub> | $c_-$ | 1.0 | $\text{min}^{-1}$ |
| Binding of Plasmid<sub>k</sub> and Protein<sub>u</sub> | $c_+$ | 1.0 | $\text{nM}^{-1} \cdot \text{min}^{-1}$ |
| Production rate of Protein<sub>x</sub> (per plasmid) | $\beta$ | 0.097 | $\text{min}^{-1}$ |
| Effective production rate of Protein<sub>u</sub> | $\alpha$ | 4.85 | $\text{nM} \cdot \text{min}^{-1}$ |
| Degradation rate of proteins | $\gamma$ | 0.035 | $\text{min}^{-1}$ |

Figure 2(a) and (b) show the changes in the total concentration of the plasmid, *k*(*t*) + *k*_*b*_(*t*), and the concentration of Protein_x_, *x*(*t*), respectively. The simulation results indicate that when *k*(*t*) + *k*_*b*_(*t*) changes in time, the synthesis of Protein_x_, *x*(*t*), changes in response to the plasmid copy number *k*(*t*). In particular, the concentration *x*(*t*) at steady state increases linearly with *k*(*t*) + *k*_*b*_(*t*). These results suggest that adjusting the plasmid copy number provides an effective and precise means of tuning the circuit gain. In the next section, we experimentally implement a circuit in which replication initiator protein can be tuned by widely used regulatory proteins and chemical inducers, providing an alternative option to tuning biocircuits in addition to the ones in the literature.^9^

**Figure 2:**
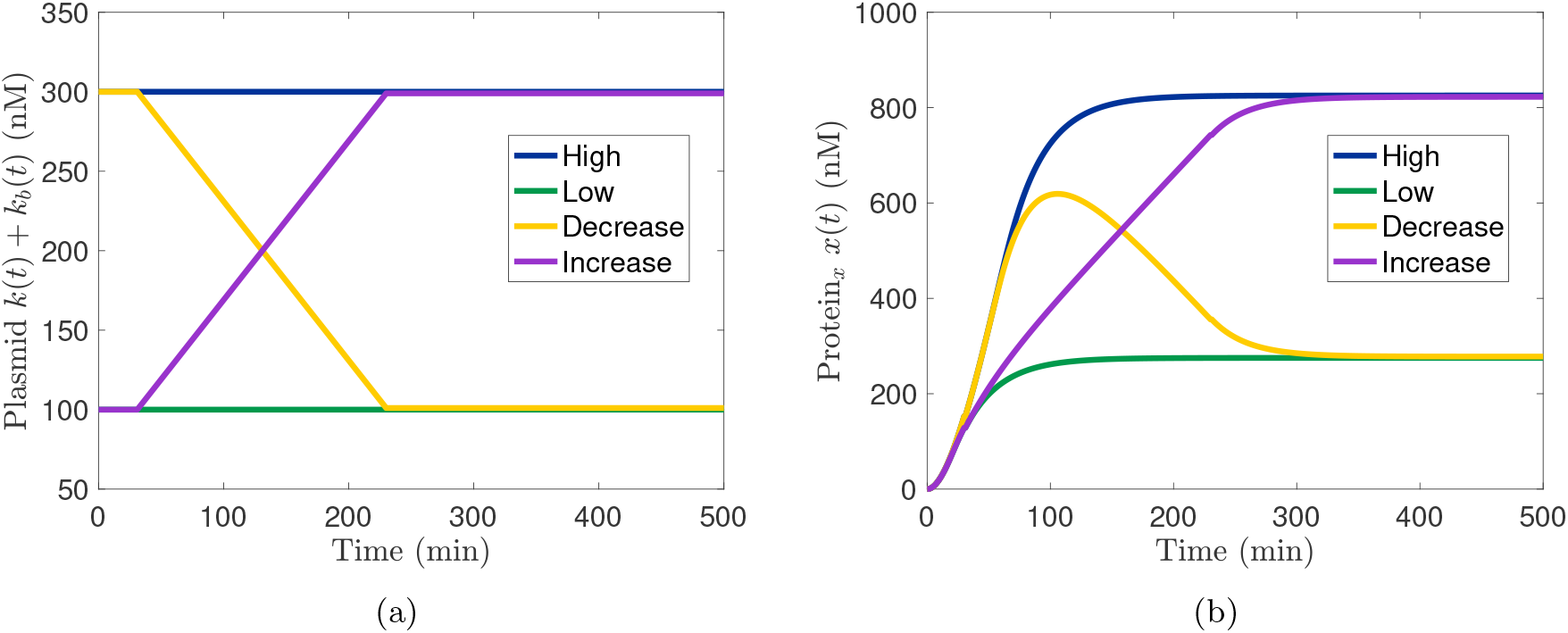
Simulated time-course of (a) Total plasmid concentration *k*(*t*) + *k*_*b*_(*t*) and (b) output protein concentration *x*(*t*). Lengeds indicate *k*(*t*) at *t* = 30 and *t* = 230, respectively. The output protein concentration is proportional to the plasmid concentration, demonstrating that the circuit gain can be modulated through plasmid copy number control.

### 2.3 Modulation of plasmid copy number by chemical inducer

We designed the circuit to allow external control of plasmid copy number by utilizing commonly used regulatory proteins and chemical inducers. Specifically, we replaced the promoter of the gene encoding the replication initiator protein RepA of pSC101 plasmid with a synthetic hybrid promoter J23110-lacO1 to enable transcriptional regulation of RepA by LacI. Based on the observation in literature,^14^ we used a variant of RepA that carried four mutations to increase the dynamic range of the copy numbers. The overall schematic diagram of the circuit is shown in Fig. 3, where the RepA variant RepA^*^ is regulated by the repressor protein LacI, which is, in turn, activated by LuxR in the presence of AHL (acyl-homoserine lactone). A gene cassette that constitutively expresses superfolder green fluorescent protein (sfGFP) was introduced to monitor its copy number through the fluorescence measurement. This circuit allows the plasmid copy number to be regulated by two common inducers, AHL and Isopropyl *β*-d-1-thiogalactopyranoside (IPTG).

**Figure 3:**
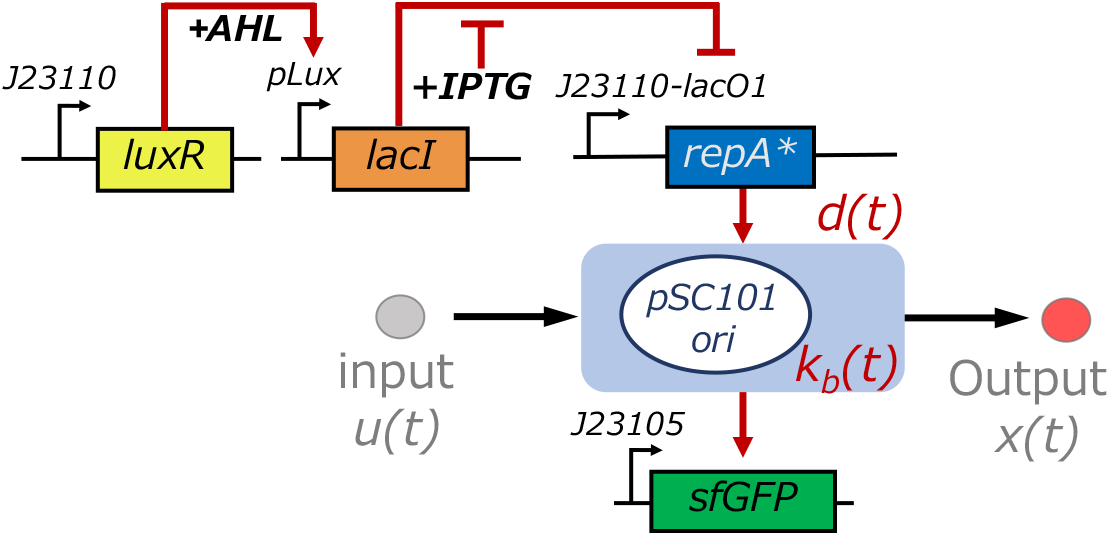
Schematic diagram of the genetic circuit for plasmid copy number control. The circuit enables external regulation of RepA^*^, a variant of replication initiator protein of pSC101 plasmid, and controls the sensitivity of the output *x*(*t*) in response to the input *u*(*t*)

To verify changes in the plasmid copy number by AHL, we performed titration experiments in *E. coli* DH5*α* harboring the circuit plasmid. Specifically, Figures 4(a) and (b) show the time-series data and the snapshot distribution at 7 hours after induction for constitutively expressed sfGFP, respectively. The copy number of the plasmid was also measured by quantitative PCR (qPCR) using the cell population collected at 4 hours after induction as shown in Fig. 4(c). As expected, increasing the AHL concentration resulted in the decrease in fluorescence intensity, indicating that the plasmid copy number was reduced in response to AHL induction. These measurements as well as Fig. 4(c) are consistent with our design expectation that the plasmid copy number and the rate of the resulting gene expression can be effectively regulated by the commonly used inducer (external input) through the modulation of RepA^*^.

**Figure 4:**
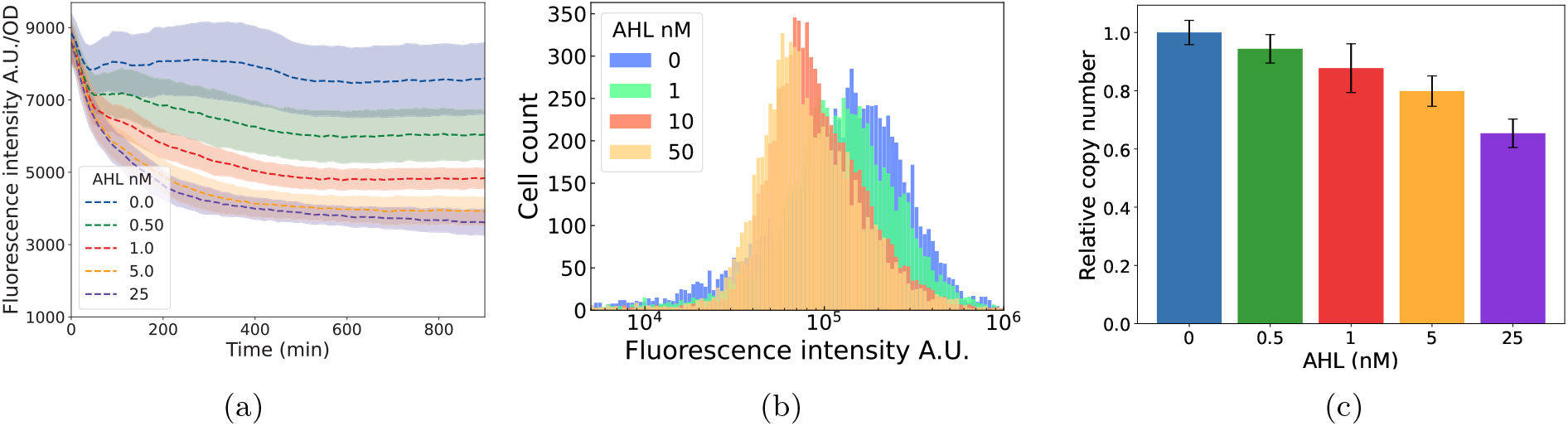
Experimental verification of copy number changes (a) time-course of sfGFP fluorescence using a plate reader. The shaded area indicates the standard error of *N* = 3 samples. (b) Snapshot of fluorescence intensity at 7 hours after induction, measured by flow cytometry (c) Normalized plasmid copy number measured by quantitative PCR. The error bar indicates standard error.

### 2.4 Modulation of plasmid copy number by cell-cell communication

Finally, we implemented a two-strain co-culture system consisting of sender and receiver cells to demonstrate copy number control by an upstream circuit via cell-cell communication (Fig. 5(a)). The upstream “controller” (sender) strain produced the quorum sensing molecule AHL by constitutively expressing the AHL synthase gene *luxI* under the control of either a strong (J23114) or weak (J23117) promoter. The strain described in Section 2.3 was used as the receiver.

**Figure 5:**
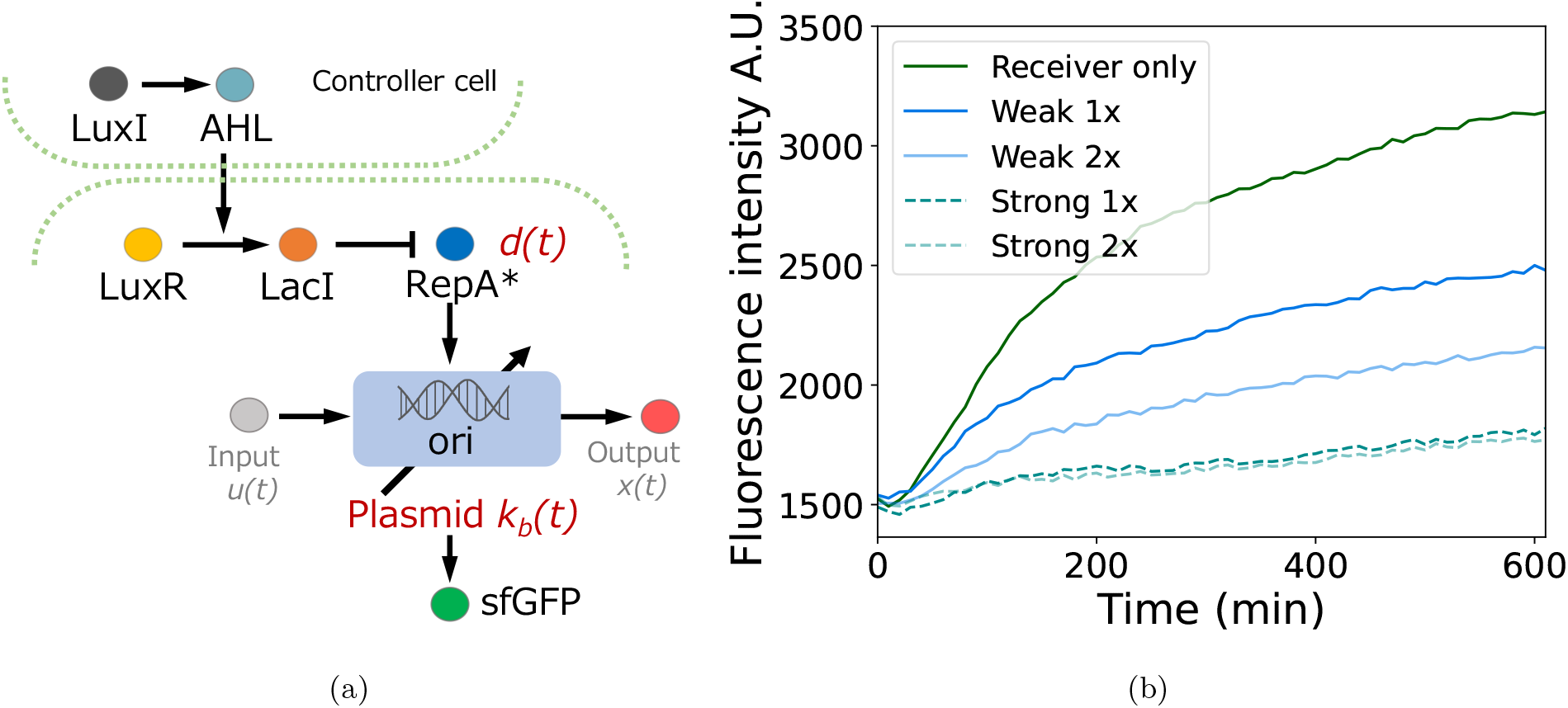
(a) Schematic figure of the genetic circuit regulated by quorum sensing molecule. (b)sfGFP fluorescence intensity under different promoter strengths and initial cell densities (1x and 2x indicate initial cell densities of OD600=0.15 and 0.30, respectively)

To evaluate the effect of the upstream signal strength, co-culture experiments were performed for each sender-receiver pair at two different initial optical densities (OD600 = 0.15 and 0.30), and fluorescence of sfGFP in the receiver cells was quantified (Figure 5(b)). In all co-culture conditions, the receiver strain exhibited reduced fluorescence level compared to the monoculture condition, reflecting repression of RepA expression via AHL-mediated signaling from the sender strain. Moreover, a graded response of the fluorescence was observed corresponding to the strength of the promoter driving *luxI* and the initial cell density. These results suggest that plasmid copy number can be effectively regulated by the output of the upstream circuit and the cell density, enabling circuit gain to be tuned through upstream control element.

## 3 Conclusion

We have proposed a genetic circuit capable of dynamically controlling plasmid copy number as a means to modulate circuit gain. The circuit was designed to operate without incorporating feedback regulation in the expression of the replication initiator protein, allowing for more predictable and analyzable system behavior. The copy-number-based gain tuning was illustrated by a mathematical model and was demonstrated through computational simulations, which highlighted how changes in plasmid abundance affect the sensitivity of the system’s input-output relation. Experimental validation confirmed that the copy number can be adjusted by external inputs. Furthermore, we demonstrated cell-to-cell communication-based regulation of plasmid copy number, enabling coordinated gain control across different strains and thereby highlighting the versatility of the approach. These results demonstrate the utility of plasmid copy number control as a practical strategy for tuning the gain of synthetic biocircuits, offering a new control handle for the design and implementation of genetic circuits.

## Material and Methods

### Simulation

All simulations in Fig. 2 were performed with MATLAB2022 using the ode45 solver with default options.

### Plasmids and Strains

The circuit plasmid in Fig. 3, which we refer to as pHNvar5, was derived from a previously reported pSC101 variant plasmid pTHSSe_47,^14^ which was a gift from Christopher Voigt (Addgene plasmid #109244). The plasmid backbone was first modified by Golden Gate assembly to replace the original promoter and ribosome binding site (RBS) upstream of *repA* gene with the synthetic constitutive promoter J23110 from the Anderson promoter library^17^ followed by a lacO1 operator and the UTR1 RBS from the literature^18^ to enable tunable expression. The *repA* variant of pSC101 origin, which originally contained previously reported two mutations R43W(A127T) and T304I(C911T), also carried two additional mutations L174S(T521C) and G259R(G775T)in the final plasmid. Next, two functional gene cassettes were constructed separately, each consisting of a promoter, gene of interest, and terminator: (i) *lacI* gene was placed under the control of the pLux promoter followed by BBa_J34803 RBS in the registry of iGem standard biological parts, and (ii) *luxR* gene was placed under the control of J23110 promoter^17^ with UTR1 RBS.^18^ The *lacI* cassette was assembled into the modified plasmid using Golden Gate assembly, and subsequently, the *luxR* cassette was inserted using Gibson assembly. The final plasmid was transformed into E. coli DH5*α* (Takara Bio Inc.) via electroporation. Transformed cells were cultured at 37°C for 16 hours and stored as glycerol stocks at −80°C.

The plasmids in Fig. 5(a), which we refer to as pHN037 and pHN038, carrying the *luxI* gene under the control of different constitutive promoters, were constructed by placing *luxI* gene under the control of either J23117 (strong) or J23114 (weak) promoter from Anderson promoter library^17^ followed by UTR1 from the literature^18^ and assembling into pBR322 plasmid backbone using Golden Gate assembly. The plasmids were transformed into E. coli DH5*α* (Takara Bio Inc.) via electroporation, incubated at 37°C for 16 hours, and stored as glycerol stocks at −80°C.

### Measurement of copy number changes by chemical induction

The data shown in Fig. 4(a)–(c) were obtained from independent cultures. In all experiments, overnight cultures were prepared by inoculating the glycerol stock into 5mL of LB medium with 100 µg/mL of Ampicillin and incubating overnight at 37°C with shaking at 225 rpm (BR-23FP, Taitec).

For the plate reader measurement (Fig. 4(a)), the overnight cultures were diluted with 5 mL of LB medium with 100 µg/mL of Ampicillin to an OD600 of 0.30 and then were incubated at 37°C with shaking speed at 225 rpm, to obtain pre-culture for the experiment. After 2 hours of incubation, the pre-culture was removed from the bioshaker, and the cell density (OD600) was measured and adjusted to 0.30 if necessary. At this stage, AHL was added to the culture, and 200 µL aliquots were transferred into the wells of a black-wall, clear bottom, flat-bottom 96-well plate (CELLSTAR, Greiner Bio-One). Fluorescence intensity and OD600 of each well was measured at every 10 minutes using a plate reader (Synergy HTX, Biotek) pre-warmed to 37°C with the 485/528 nm excitation/emission filters and a gain setting of 40.

For the snapshot measurement in Fig. 4(b), the pre-culture was prepared in the same manner as described above. After 2.5 hours of incubation, the pre-culture was removed from the bioshaker, and OD600 was adjusted to 0.30 by diluting with LB medium to a final volume of 5 mL. Then, AHL was added to each culture,. and the cultures were further incubated in the bioshaker at 37°C with shaking speed at 225 rpm. After 7 hours of incubation, each culture was loaded into a flow cytometer (BD Accuri C6, BD Biosciences) to measure sfGFP fluorescence intensity using the FL1A channel.

For the qPCR experiment in Fig. 4(c), the cultures were prepared and induced by AHL in the same manner as the snapshot experiment described above. Each culture was incubated in the bioshaker at 37°C with shaking speed at 225 rpm. After 4 hours of incubation, 3 aliquots were taken from the culture tubes and adjusted to OD600 of 1.0. 10 µL of each aliquot was then mixed with 90 µL of ultrapure water, and the mixture was boiled at 95°C for 20 minutes in a thermal cycler (T100 Thermal Cycler, Bio-Rad) to lyse the E. coli cells. The resulting lysates were used for qPCR. Reactions were prepared with PowerTrack SYBR Green Master Mix (ThermoFisher Scientific) and previously reported primers (BLA F/R for the plasmid-borne ampicillin resistance gene and DXS F/R for the chromosomal *dxs* gene, present at 1 copy per genome).^14^ qPCR was performed on AriaMx Real-time PCR System (Agilent) using the following protocol: 95°C for 2 min; 40 cycles of 95°C for 15 s, 60°C for 1 min. Relative plasmid copy number was calculated using the ΔΔCt method with a plasmid containing a single ampicillin resistance gene and *dxs* as the calibrator assuming an efficiency of 100%.

### Measurement of copy number changes by cell-cell communication

*E. coli* DH5*α* strains harboring pHNvar5, pHN037, and pHN038 were inoculated from glycerol stocks, and pre-cultures were prepared using the same protocol as the plate reader measurement described above. After removing the pre-culture tubes from the bio-shaker, OD600 was adjusted to 0.30 for the sender strain harboring pHNvar5 and either 0.60 or 1.20 for the receiver strains harboring pHN037 or pHN038 using LB medium.

For each condition, 150 µL of the sender strain was mixed with 50 µL of the receiver strain. The co-cultures were transferred to a black-wall, clear bottom, flat-bottom 96-well plate (CELLSTAR, Greiner Bio-One). The fluorescence intensity and OD600 were measured using the plate reader, pre-warmed to 37°C, at every 10 minutes using 485/528 nm excitation/emission filters and a gain setting of 40.

## Acknowledgments

This work was supported in part by JSPS KAKENHI Grant Number JP23H01355, the Ishii-Ishibashi Fund (The Keio University Grant for Early Career Researchers), and JST-Mirai Program Grant Number JPMJMI22G6.

